# Gigaxonin, mutated in Giant Axonal Neuropathy, interacts with TDP-43 and other RNA binding proteins

**DOI:** 10.1101/2024.09.03.611033

**Authors:** Cassandra L. Phillips, Maryam Faridounnia, Rachel A. Battaglia, Baggio A. Evangelista, Todd J. Cohen, Puneet Opal, Thomas W. Bouldin, Diane Armao, Natasha T. Snider

## Abstract

Giant Axonal Neuropathy (GAN) is a neurodegenerative disease caused by loss-of-function mutations in the *KLHL16* gene, encoding the cytoskeleton regulator gigaxonin. In the absence of functional gigaxonin, intermediate filament (IF) proteins accumulate in neurons and other cell types due to impaired turnover and transport. GAN neurons exhibit distended, swollen axons and distal axonal degeneration, but the mechanisms behind this selective neuronal vulnerability are unknown. Our objective was to identify novel gigaxonin interactors pertinent to GAN neurons. Unbiased proteomics revealed a statistically significant predominance of RNA-binding proteins (RBPs) within the soluble gigaxonin interactome and among differentially-expressed proteins in iPSC-neuron progenitors from a patient with classic GAN. Among the identified RBPs was TAR DNA-binding protein 43 (TDP-43), which associated with the gigaxonin protein and its mRNA transcript. TDP-43 co-localized within large axonal neurofilament IFs aggregates in iPSC-motor neurons derived from a GAN patient with the ‘axonal CMT-plus’ disease phenotype. Our results implicate RBP dysfunction as a potential underappreciated contributor to GAN-related neurodegeneration.

**Summary:** This work reveals that the neurodegeneration-associated protein and cytoskeleton regulator gigaxonin and its mRNA associate with numerous RNA binding proteins. These findings shift understanding of normal gigaxonin function and provide insights into how disease-causing mutations in the gigaxonin-encoding gene (*KLHL16*) may ignite a pathogenic cascade in neurons.

## Introduction

Giant axonal neuropathy (GAN) is a pediatric monogenic neurodegenerative disease that affects the peripheral nervous system (PNS) and the central nervous system (CNS) (Johnson-Kerner et al., 2014). Neurons degenerate due to loss-of-function mutations in the *KLHL16* gene (also known as *GAN*), which encodes the protein gigaxonin (Bomont et al., 2000). Gigaxonin regulates the transport (Renganathan et al., 2023) and degradation (Mahammad et al., 2013) of cytoskeletal intermediate filament (IF) proteins and, in its absence, IFs accumulate in many cell types (Mahammad et al., 2013; Opal and Goldman, 2013). The clinically debilitating effects of GAN are due to the preferential and pronounced involvement of neurons. The pathologic hallmark of GAN is the presence of ‘giant’ axons distended by focal, densely packed neuronal IFs, including neurofilaments (NFs) and peripherin (Berg et al., 1972; Johnson-Kerner et al., 2015a). It is not known why neurons are the most vulnerable cell type in GAN and if the accumulation of IF proteins is central to the GAN-related neurodegeneration.

In GAN patients, early developmental milestones are normal (Bharucha-Goebel et al., 2021). Classic GAN, which involves both the CNS and PNS, is heralded by clumsiness of gait at 2-3 years of age and loss of independent ambulation in the first decade of life. A milder form of GAN that has an ‘axonal Charcot-Marie-Tooth disease plus’ (CMT-plus) phenotype and fewer of the systemic and CNS manifestations of classic GAN has also been recognized and has a mean age of onset of 5-6 years. Advances in next generation sequencing may enable earlier diagnoses of milder GAN cases, reported to carry at least one missense variant in the *KLHL16/GAN* gene. Currently, there are more than 400 variants of uncertain significance reported for *KLHL16/GAN* in ClinVar, and >75% of these variants are missense mutations (Landrum et al., 2016).

Here we sought to identify gigaxonin interactors that have direct relevance to the cell biology of GAN neurons, particularly during the early stages of neuronal differentiation. Our approach combined unbiased mass spectrometry-based proteomics to identify new gigaxonin interactors, unbiased profiling of differentially expressed proteins in GAN neural progenitor cells (NPCs), and validation studies in induced pluripotent stem cell (iPSC)-motor neurons (MNs) of GAN patients. This led us to the discovery of numerous RNA-binding proteins (RBPs) as novel and underappreciated interactors of gigaxonin that are also altered in GAN patient cells. Importantly, we discovered that a subset of RBPs, including TDP-43, associate with both the gigaxonin protein and its mRNA. Collectively, our results reveal complex cellular disease mechanisms that pave the way for examining altered RNA biology as a driver in GAN neurodegeneration, and move beyond the current focus on the IF cytoskeleton.

## Results

### RBPs predominate the soluble gigaxonin interactome

IF proteins are highly abundant and frequently identified in mass spectrometry analyses of total cell and tissue lysates. While this is advantageous for identifying specific IFs within complex mixtures, it can also mask the presence of lower abundance proteins. IF proteins are solubilized in SDS detergent-containing lysis buffer (like RIPA), but are sparingly soluble in lysis buffer containing the non-ionic TritonX-100 detergent (Snider and Omary, 2016; Yang et al., 2022). We reasoned that using a non-ionic extraction method would help us uncover less abundant, but biologically relevant, gigaxonin interactors. We over-expressed GFP-tagged human gigaxonin (GFP-Gig), or GFP vector control, in HEK293 cells and isolated the TritonX-100 soluble fraction. This enabled us to compare our results to a previous proteomic analysis (Johnson-Kerner et al., 2015b) of gigaxonin interactors from RIPA-soluble lysates (**Fig. 1A**). We analyzed the high confidence gigaxonin interactors identified in the prior study (Johnson-Kerner et al., 2015b) using the Gene Ontology (GO):Molecular Function tool in DAVID (Sherman et al., 2022). As expected, ‘structural constituent of the cytoskeleton’ was the most significant category of proteins that interact with gigaxonin in the RIPA-soluble lysates (**Fig. 1B**). However, there were also other significant gigaxonin interactors, including RNA-binding proteins (RBPs), protein folding chaperones, ATP-binding proteins, and structural constituents of ribosomes (**Fig. 1B**). Upon validating GFP-Gig over-expression in HEK293 cells (**Fig. 1C**), we analyzed the TritonX-100-soluble proteins present in a GFP pulldown of untransfected, GFP vector control and GFP-Gig expressing cells. We identified 545 proteins with >2 log-fold-change (LFC) and reaching statistical significance (p<0.05) in the GFP-Gig pulldown compared to untransfected and GFP vector-transfected cells (**Supplemental Table 1**). As expected, IF proteins, including vimentin, keratins 8/18, and lamins B1/B2 were among the gigaxonin interactors, but they were not the most highly enriched targets (**Fig. 1D**). The proteins with the highest log fold change (LFC) and significance were various RBPs (e.g. HNRNPF, PABPC1, ZC3HAV1); ribosomal proteins (e.g. RPL6, RPL7, RPL15); folding factors (PFDN5, NUDCD3); and ubiquitin ligases (STUB1, ARIH1, TRIM38, HUWE1) (**Fig. 1D**). Strikingly, RBPs as a group comprised 40% of all gigaxonin interactors and were the highest significance category (P=7.4E-102) based on the GO:Molecular Function analysis in DAVID (**Fig. 1E**). The 217 RBPs that we identified in the GFP-Gig pulldown (**Supplemental Table 2**) are involved in cytoplasmic translation and splicing, ribosome biogenesis, translation initiation and mRNA stabilization (**Fig. 1F**). Therefore, our modified proteomics method aimed at uncovering soluble gigaxonin interactors revealed RBPs as a major category.

**Figure 1.**
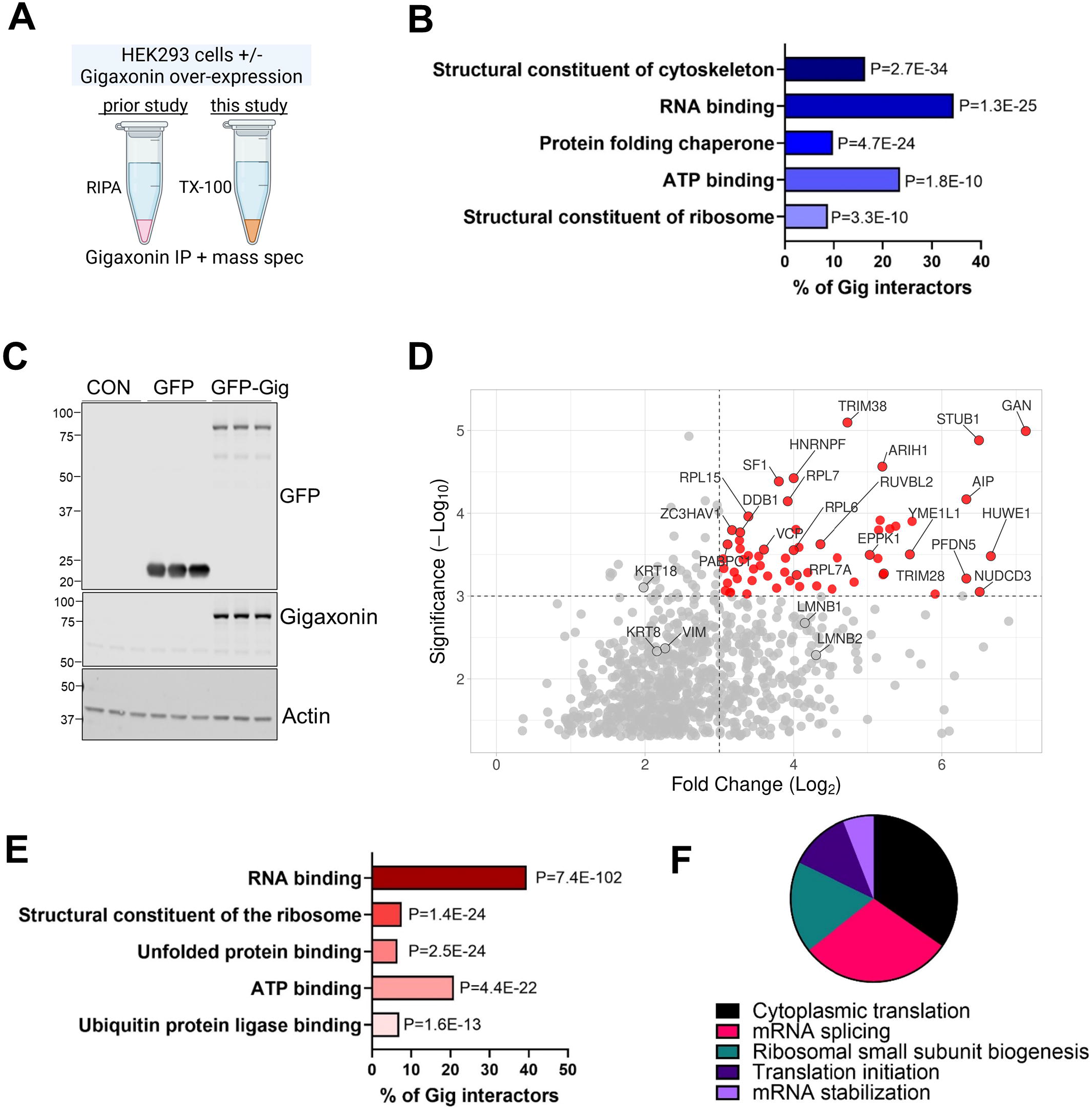
RBPs predominate the soluble gigaxonin interactome. **A**. Schematic of our current method compared to a prior published proteomic study (Johnson-Kerner et al., 2015b). **B**. Classification of gigaxonin interactors (identified in the published dataset) based on known molecular function. Clustering and P-values were obtained using the DAVID bioinformatics resource. **C**. GFP and gigaxonin immunoblots of HEK293 cell lysates transfected with Lipofectamine-3000 alone (lanes 1-3), GFP vector alone (lanes 4-6), or GFP-gigaxonin (lanes 7-9). Actin serves as loading control. **D**. Volcano plot of 545 proteins enriched in the GFP-gigaxonin (GFP-Gig) pulldown. Highlighted in red are proteins with the highest log fold change (LFC) and statistical significance. **E**. Major functional categories in the GFP-Gig pulldown. Stratification was performed by Gene Ontology (GO)-Molecular Function (MF) using DAVID. **F**. Major biological processes associated with the RNA-binding proteins (RBPs) identified in the gigaxonin pulldown. Stratification was performed by Gene Ontology (GO)-Biological Process (BP) using DAVID.

### RBPs are the top category of differentially expressed proteins in GAN patient NPCs

To test the biological significance of these results in a GAN-relevant cellular context, we used patient-derived neural progenitor cells (NPCs) and corresponding isogenic controls. The GAN and control iPSCs were derived from a patient with classic GAN, carrying the homozygous gigaxonin point mutation G332R, and were described recently (Battaglia et al., 2023). We compared 3 independent clones of each line (unedited and revertant). Gigaxonin protein was extremely low in the unedited GAN mutant cells (consistent with prior studies (Battaglia et al., 2023; Boizot et al., 2014)), but was restored in the isogenic control cells (**Fig. 2A**). Both GAN and control NPCs expressed the NPC marker βIII-tubulin (**Fig. 2B**). Typically, NF expression is low in the early stages of neuronal differentiation and NFs become significantly upregulated as neurons mature (Pachter and Liem, 1984), but NF expression dynamics during development and in adulthood have not been examined in the context of GAN. In the unedited GAN patient NPCs, we observed aggregates of phosphorylated NF-heavy (pNF-H), a known marker of neurodegeneration (Snider and Omary, 2014; Yuan and Nixon, 2021; Yuan et al., 2012). This led us to ask if there were early proteome-wide changes in GAN. Using unbiased proteomics, we identified 297 proteins with significant differential expression (p<0.05) between GAN and control NPCs (**Supplemental Table 3**). The most upregulated protein in GAN NPCs was the serine/threonine-protein phosphatase 4 catalytic subunit (PPP4C), which has many reported roles in microtubule organization (Helps et al., 1998; Toyo-oka et al., 2008), maturation of spliceosomal small nuclear ribonucleoproteins (Carnegie et al., 2003), and neuronal differentiation (Lyu et al., 2013). NFs and α-internexin (INA) were also among the most upregulated proteins (**Fig. 2C**). Several metabolic enzymes (e.g. methionine transferase [MTR] and Acyl-CoA 6-desaturase [FADS2]) and mitochondrial proteins (e.g. misato homolog 1 [MSTO1] and NADP transhydrogenase [NNT]) were among the most downregulated – consistent with the known mitochondrial abnormalities in GAN (Israeli et al., 2016). However, collectively, RBPs were the most over-represented category of differentially-expressed proteins with the highest statistical significance (P=5.6E-15) (**Fig. 2D**). Combined with the soluble gigaxonin interactome data, these results suggest that RBP levels and/or their functions may potentially be altered in the presence of GAN disease-causing mutations.

**Figure 2.**
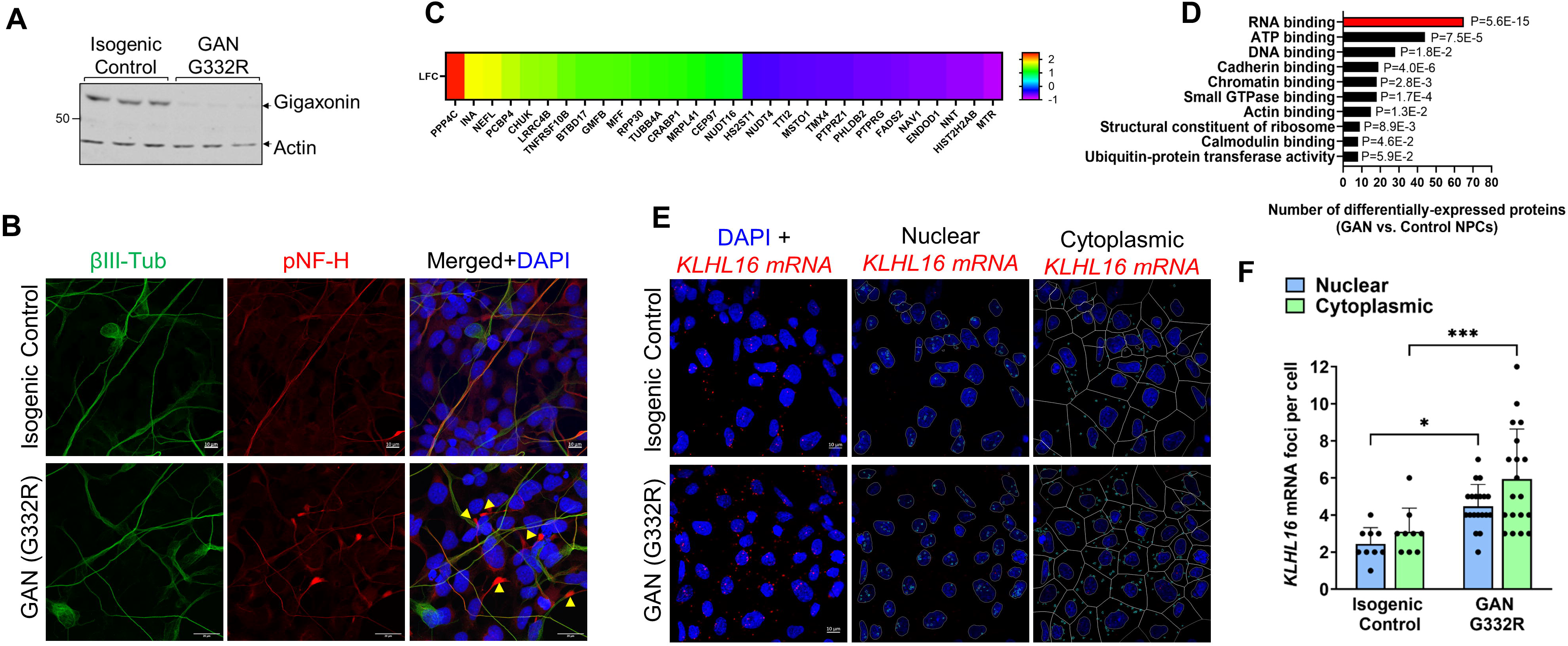
RBPs are the major category of differentially expressed proteins in GAN NPCs. **A**. Immunoblot comparison of gigaxonin expression in three independent clones from GAN mutant (G332R) and three clones of the corresponding isogenic control NPCs. Actin serves as loading control. **B**. Immunofluorescence staining of GAN and control NPCs showing expression of βIII-tubulin (marker of NPCs; green) along with phosphorylated NF-heavy (pNF-H; red) and DAPI (blue). **C**. Heat map of the top most upregulated and downregulated proteins (denoted by their corresponding gene names) in GAN compared to control NPCs. **D**. Major categories of differentially-expressed proteins identified by mass spectrometry analysis of GAN and control NPCs. Stratification was performed by Gene Ontology (GO)-Molecular Function (MF) using DAVID. **E**. Immunofluorescence analysis of *KLHL16* mRNA (red) and DAPI nuclear staining (blue) in control and GAN NPCs. The cytoplasmic and nuclear *KLHL16* puncta were analyzed using Cell Profiler. **F**. Quantification of nuclear (blue) and cytoplasmic (green) *KLHL16* mRNA puncta in control and GAN NPCs. ^*^p<0.05; ^***^p<0.001. Two-way ANOVA.

### Gigaxonin mRNA (KLHL16) is elevated in the cytoplasm and nuclei of GAN patient NPCs

Currently it is not clear whether the loss of gigaxonin protein in GAN cells is due to altered *KLHL16* mRNA processing, protein instability, or a combination of factors specific to the particular genetic mutation. We observed previously that the human *KLHL16* mRNA (NM_022041) has an exceptionally long 3′ untranslated (UTR) region and accumulates in the nuclei of GAN patient fibroblasts with vimentin IF aggregates (Battaglia et al., 2023). This finding suggests potential interplay between IF pathology and *KLHL16* mRNA processing in GAN. Therefore, we asked if the *KLHL16* mRNA is altered in GAN compared to control NPCs. While there was no major difference in terms of its cytoplasmic vs. nuclear distribution, we detected significantly more *KLHL16* mRNA puncta in the GAN cells compared to the isogenic controls (**Fig. 2E-F**). These observations raise the possibility of potential altered interactions between the *KLHL16* mRNA and regulatory RBPs in GAN cells. Exactly how the gigaxonin coding mutation (G332R), present in our classic GAN cell model, could exert influence over the regulation of its transcript is not clear. However, it is plausible that the presence of the long 3′ UTR region in the *KLHL16* mRNA may play a role.

### Gigaxonin protein and its mRNA share RBP interactors, including TDP-43

In general, 3′ UTRs bind to RBPs to regulate mRNA processing and, in some cases, RBPs can bind to both the 3′ UTR of the mRNA and its protein product (Gallicchio et al., 2023). Therefore, we used publically available global RNA-binding data from enhanced crosslinking and immunoprecipitation (eCLIP) studies (Van Nostrand et al., 2016) to ask if gigaxonin and its mRNA share RBP interactors. Based on the eCLIP data, which were obtained through RNAct (Lang et al., 2019) and shown in **Fig. 3A**, human *KLHL16* associates with 18 different RBPs. We identified 5 of the *KLHL16*-interacting RBPs (abbreviated by gene name: EFTUD2, PCBP2, YBX3, TARDBP, and KHSRP) as interactors of gigaxonin in our pulldown assay (**Supplemental Table 1**). Two of the overlapping RBPs, the poly(rC)-binding protein 2 (PCBP2; also known as hnRNP E2) and TDP-43 (TARDBP) have defined motifs to which they bind (Dominguez et al., 2018). Therefore, we searched for the presence of such motifs on the *KLHL16* mRNA using the RBPMap tool (Paz et al., 2014). As shown in **Fig. 3B-C**, there are numerous binding motifs for both PCBP2 and TDP-43, with the highest confidence motifs localized to the 3′ UTR region of the *KLHL16* mRNA, between nucleotides ∼2000-15000. Next, we validated our proteomic results, demonstrating that these two RBPs co-immunoprecipitate with gigaxonin (**Fig. 3D-E**). Both PCBP2 and TDP-43 are known to co-localize within pathological aggregates in neurodegenerative disease (Davidson et al., 2017; Kattuah et al., 2019). This novel association between gigaxonin and RBPs warrants further investigation, as it alludes to additional functions for gigaxonin beyond its role as a cytoskeleton regulator. Collectively, our data reveal that RBPs known to be of critical importance to neuronal homeostasis, particularly TDP-43, associate with both the gigaxonin mRNA and its protein product, and that this may be altered by GAN-causing mutations.

**Figure 3.**
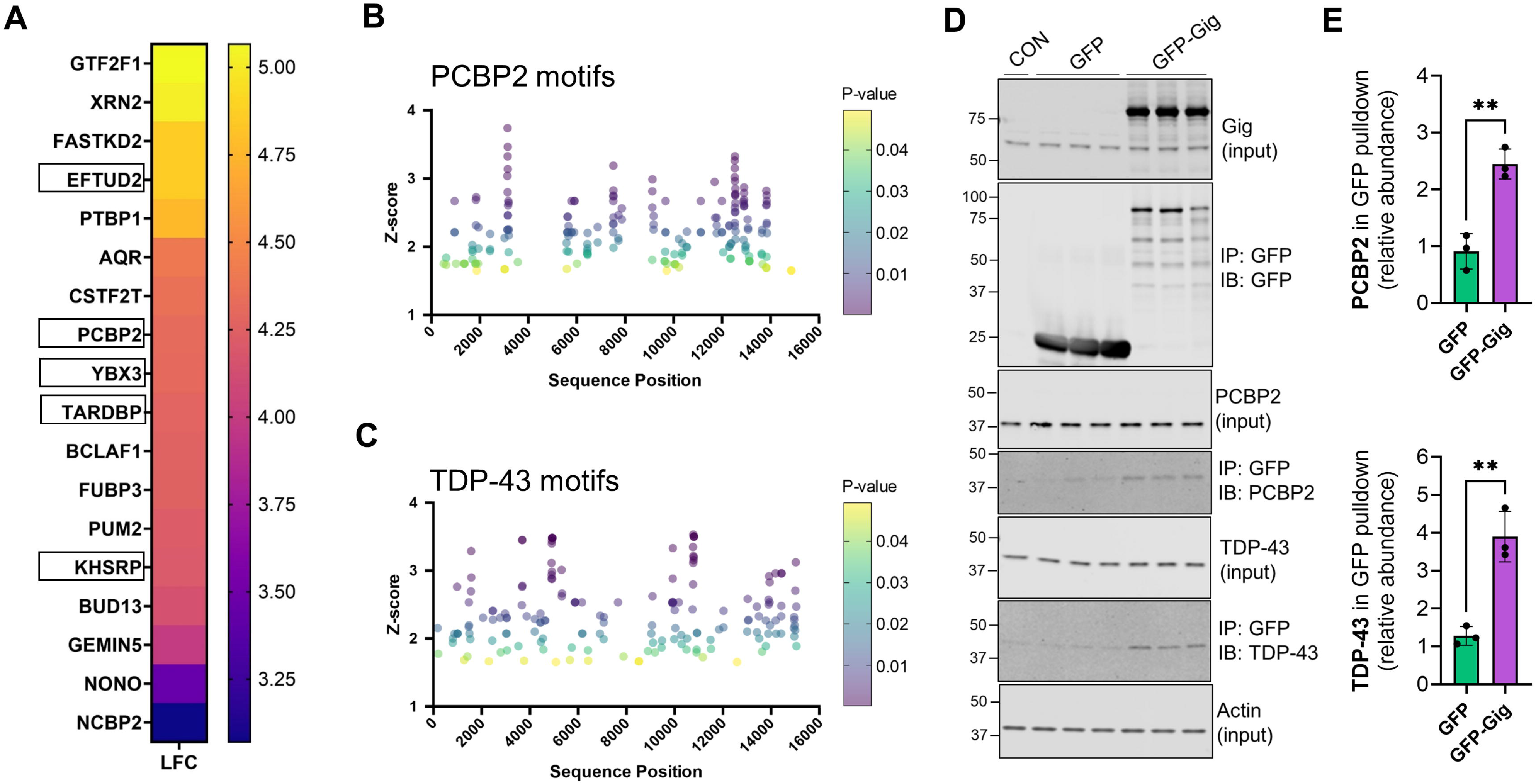
Gigaxonin protein and its mRNA share RBP interactors. **A**. Heat map showing fold enrichment of different RBPs for human *KLHL16* mRNA, obtained from publicly available eCLIP data via RNAct. Boxed are five RBPs that also interact with gigaxonin protein, based on our current proteomics study (Supplemental Table 1). **B-C**. Plots of z-scores (y-axis; indicating motif confidence) of PCBP2 (**B**) and TDP-43 (**C**) binding motifs along the *KLHL16* mRNA sequence position (x-axis). The color of the dots indicates the p-value. **D**. Validation of the mass spec results showing that PCBP2 and TDP-43 co-immunoprecipitate with gigaxonin. **E**. Quantification of PCBP2 and TDP-43 in the GFP-Gig pulldowns.

### GAN iPSC-motor neurons display granular structures and perinuclear inclusions of IFs and TDP-43

TDP-43 is normally localized to the nucleus, but undergoes nuclear-cytoplasmic shuttling to regulate RNA stabilization, transport and splicing (Tziortzouda et al., 2021). Given its association to gigaxonin, we asked if TDP-43 is mis-localized in GAN patient neurons. We did not observe major changes in TDP-43 localization in the classic GAN patient iPSC-motor neurons (MNs) carrying the G332R mutation, compared to isogenic controls, as most of the TDP-43 protein appeared nuclear (**Fig. 4A**). However, in the isogenic control cells, *KLHL16* mRNA puncta localized along neuronal processes that also stained diffusely for cytoplasmic TDP-43 and neurofilaments (**Fig. 4A** bottom left; arrows). In contrast, in GAN neurons we observed that the *KLHL16* mRNA was primarily in and around the nuclei, and especially appeared to ‘decorate’ the large perinuclear NF aggregates (**Fig. 4A** bottom right; arrows). Although widespread cytoplasmic accumulation of TDP-43 was not observed, some GAN neurons displayed perinuclear TDP-43 aggregates that co-localized with the gigaxonin mRNA (**Fig. 4B**). Furthermore, ultrastructural examination of the perinuclear region in GAN neurons revealed the presence of ∼200nm dense granular bodies enmeshed in a thicket of IF accumulations (**Fig. 4C-D**) that mirror previous human GAN EM findings (Koch et al., 1977). The composition of the granular bodies could not be resolved into any specific components or structural pattern but appeared only as roughly spherical, non-membrane bound, electron-dense granular condensations (**Fig. 4D**; green arrowheads). Surrounding IFs (**Fig. 4D**; black arrows) were intimately associated with the dense granular bodies, and IFs could be seen to frequently gather into the density. Therefore, the human GAN iPSC-MN cell model recapitulates the distinctive ultrastructural neuronal pathologic phenotype of IF accumulations forming complexes with dense granular bodies in human GAN patients *in vivo*, as first described decades ago (Asbury et al., 1972). The origin, nature, and significance of these granular bodies remains to be determined.

**Figure 4.**
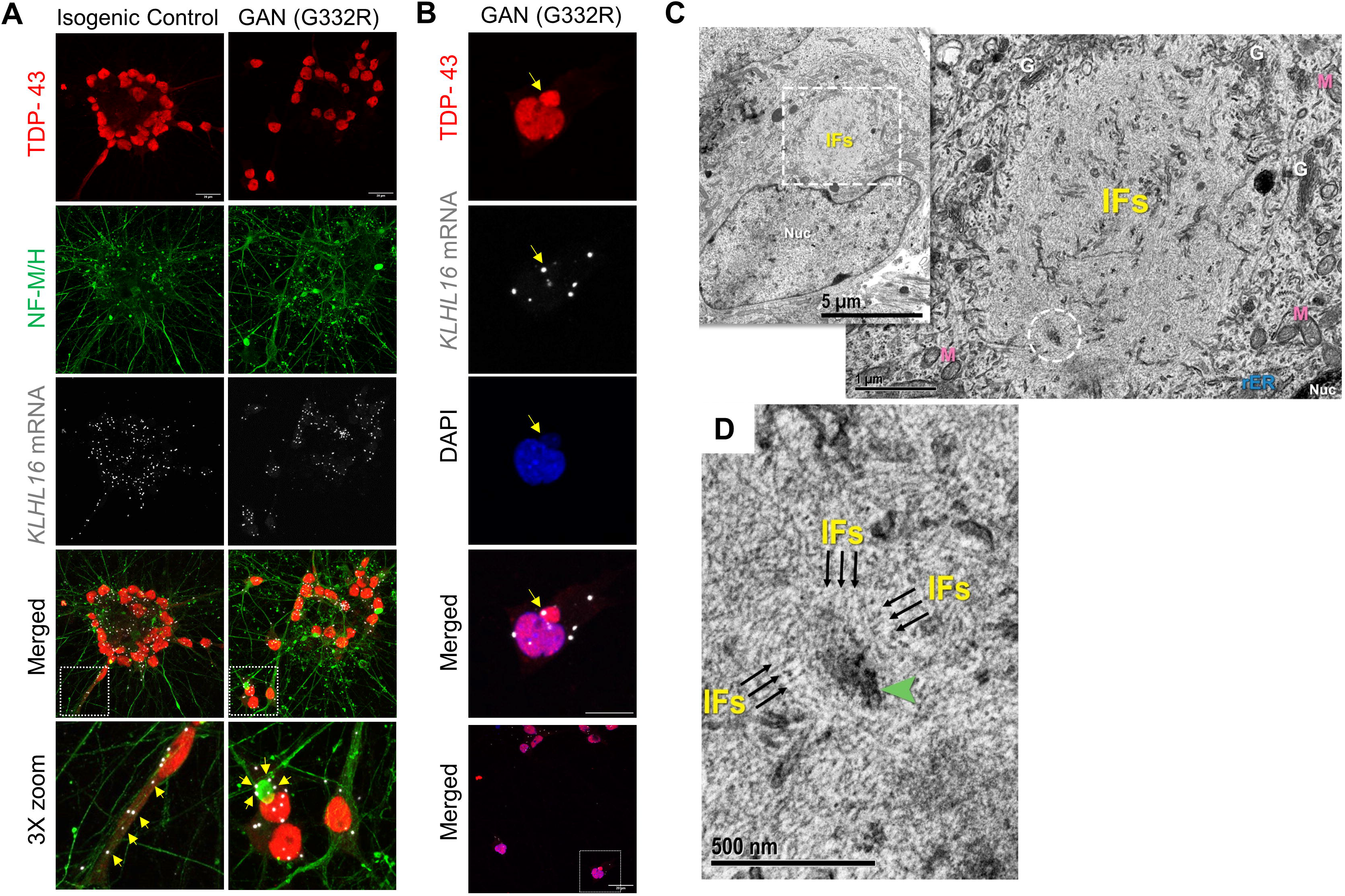
GAN iPSC-MNs display dense granular structures embedded within perinuclear IF accumulations that also co-localize with *KLHL16* mRNA. **A**. Immunofluorescence analysis of TDP-43 (red), NF-M/H (green), and *KLHL16* mRNA (white) in clusters of cell bodies from control and GAN iPSC-MNs. Scale bars=20μm. Arrows in the bottom enlarged areas point to the different localization of the *KLHL16* mRNA. **B**. Immunofluorescence staining of TDP-43 (red), *KLHL16* mRNA (white) and DAPI (blue) and in a single GAN iPSC-MNs. Arrow points to co-localized *KLHL16* with a perinuclear TDP-43 aggregate. Scale bar=10μm. The bottom image shows the selected area shown in the top three images. Scale bar=20μm. **C**. *(inset)* GAN iPSC-motor neuron (MNs) shows an irregularly contoured nucleus (Nuc) and an adjacent cytoplasmic IF aggregate (dashed box) [5kx; scale bar=5 µm]. High magnification of dashed box reveals tightly packed IFs with an entrapped electron dense granular body (dashed circle); (Nuc = nucleus, M = mitochondria, G = Golgi apparatus, rER = rough endoplasmic reticulum) [30kx; scale bar=1 µm]. **D**. Higher magnification of the dense granular body in C. shows a surrounding disorderly meshwork of IFs, many of which tie into (arrows) the non-membrane bound dense granular body (arrowhead) [80kx; scale bar=500 nm].

### TDP-43 co-accumulates within axonal NF inclusions in some GAN patient neurons

In GAN neurons, IFs accumulate both in the soma and axons. Axonal IF aggregates, a characteristic feature of GAN, are considered to be disease promoting, since they can impede axonal traffic and immobilize organelles, such as mitochondria (Israeli et al., 2016; Lowery et al., 2016). The G332R classic GAN iPSC-MNs showed limited axonal NF accumulations and were also without evidence of axonal TDP-43 accumulation when differentiated on microfluidic devices to separate the axons from the soma (**Fig. 5A**). We also examined another GAN patient line carrying a heterozygous null/R138L mutation. The latter was derived from a patient with the axonal CMT-plus phenotype, which is the milder form of GAN, with a predominantly motor and sensory polyneuropathy phenotype and little or no CNS involvement (Bharucha-Goebel et al., 2021). We found widespread axonal accumulation of TDP-43 in the null/R138L iPSC-MNs compared to the control and G332R iPSC-MNs (**Fig. 5B**). The TDP-43 co-localization was particularly strong within larger (4-5μm) NF inclusions (**Fig. 5C**). In the soma compartment of the null/R138L GAN patient cells, we found evidence of extranuclear TDP-43 localization (**Fig. 5D**). Based on the human iPSC GAN cellular models, axonal TDP-43 mislocalization may be present in some GAN patients, but this could vary depending on the type of mutation and clinical presentation of the disease. Combined, our results uncover a complex molecular mechanism in GAN involving RBP associations with both the *KLHL16* transcript and the gigaxonin protein.

**Figure 5.**
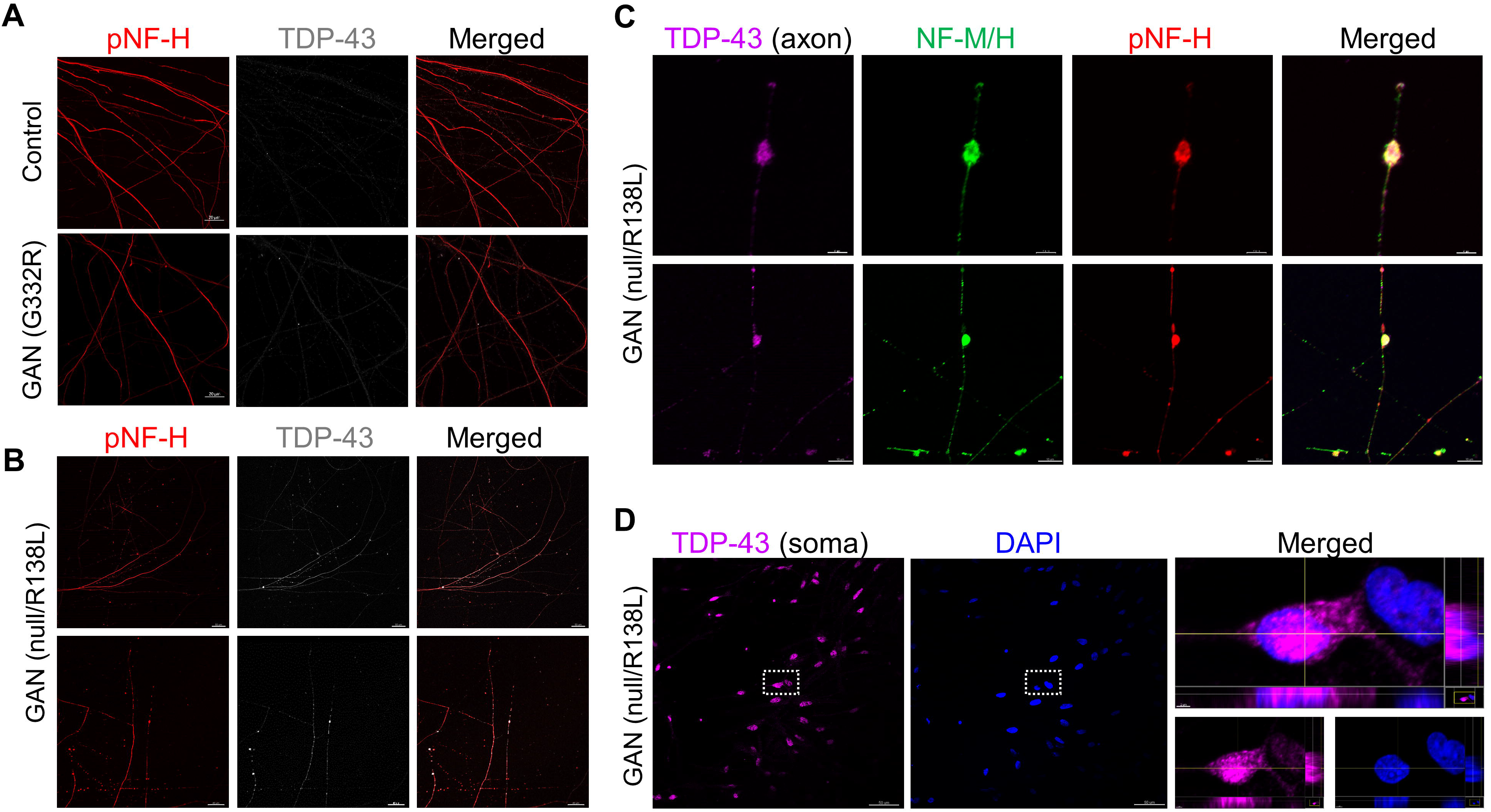
Axonal TDP-43 co-aggregates with NFs in some GAN patient-derived iPSC-MNs. **A**. Immunofluorescence co-localization of phospho-NF-H and TDP-43 in the axonal compartment of ‘classic’ GAN patient iPSC-MNs carrying the homozygous G332R mutation and corresponding isogenic control. Scale bars=20μm **B**. Immunofluorescence co-localization of phospho-NF-H and TDP-43 in the axonal compartment of ‘axonal CMT (plus)’ GAN patient iPSC-MNs carrying the compound heterozygous null/R138L mutation. Scale bars=50μm (top row); 40μm (bottom row). **C**. High magnification images of TDP-43 axonal aggregates (magenta) co-stained with total NF-M/H (green) and phospho-NF-H (red) in GAN patient iPSC-MNs carrying the null/R138L mutation. Scale bars=4μm (top row); 10μm (bottom row). **D**. Immunofluorescence co-localization of DAPI (blue) and TDP-43 (magenta) in the soma compartment of GAN patient iPSC-MNs carrying the null/R138L mutation. Scale bar=50μm (left). Images on the right show magnified view of the nuclei in the dashed box on the left. Scale bar=2μm (right).

## Discussion

Gigaxonin belongs to the Kelch-like gene family consisting of 42 members in human (Dhanoa et al., 2013). The proteins encoded by these genes share a common domain organization consisting of N-terminal BTB and BACK domains and a C-terminal Kelch domain composed of several (most often, six) Kelch motif repeats (Shi et al., 2019). The functions of most *KLHL* family members remain enigmatic, but several diseases have been attributed to mutations in these genes, pointing to important functions in the nervous system and in muscle. Aside from *KLHL16* mutations in GAN, other diseases linked to this gene family include intermediate epidermolysis bullosa simplex (EBS) with cardiomyopathy caused by *KLHL24* mutations (Yenamandra et al., 2018), early-onset distal myopathy due to *KLHL9* mutations (Cirak et al., 2010), and congenital nemaline myopathies caused by *KLHL40* (Ravenscroft et al., 2013) and *KLHL41* (Gupta et al., 2013) mutations. Functionally, the best characterized member of this family is *KLHL19*, encoding KEAP1, a substrate-specific adapter of an E3 ubiquitin ligase complex that controls NRF2 ubiquitination and plays an important role in oxidative stress and cancer. The exact function and regulation of gigaxonin remain to be elucidated, but its most widely accepted role is that of an E3 ubiquitin ligase adapter promoting the degradation of cytoskeletal IF proteins, including neurofilaments (Bomont, 2016), GFAP (Lin et al., 2016), vimentin (Mahammad et al., 2013), and epidermal keratins (Buchau et al., 2018). While some gigaxonin functions can be delineated based on related family members, like KEAP1, it is important to note that the human *KLHL16* mRNA is uniquely and exceptionally long (three times longer than other *KLHL* transcripts), and the possibility exists that gigaxonin function is highly specialized and may be uniquely regulated at the mRNA and protein level. Our work here points to a novel role of gigaxonin regulating RNA biology.

In overexpression studies, gigaxonin promotes complete clearance of both normal IFs and pathologic IF aggregates in various cellular models, including GAN patient cells (Johnson-Kerner et al., 2015a). However, normal levels of endogenous gigaxonin in the cell are very low, and therefore over-expression studies showing complete IF protein clearance have to be interpreted with some caution. Indeed, recent work shows that gigaxonin may also be involved in cellular transport, thereby regulating intermediate filament cycling and dynamics (Renganathan et al., 2023). In addition to IF proteins, other proposed substrates for gigaxonin include the microtubule-associated proteins MAP1B and MAP8 and the tubulin chaperone TBCB (Allen et al., 2005; Ding et al., 2006; Wang et al., 2005). Gigaxonin has also been implicated in autophagy through its interaction with, and proposed degradation of, the ATG16L1 protein, which is necessary for autophagosome formation (Lescouzères and Bomont, 2020; Scrivo et al., 2019). In the absence of gigaxonin, there is evidence of impaired autophagosome production due to ATG16L1 accumulation, which suggests a role for gigaxonin in regulating autophagy activity (Scrivo et al., 2019). Similar to our study, prior proteomic studies have reported interactions between gigaxonin and ribosomal proteins, but any significance of these interactions has been completely overshadowed by the strong cytoskeleton phenotypes (Johnson-Kerner et al., 2015b; Mussche et al., 2012). The work presented here has identified unexpected gigaxonin-RBPs interactions, not only at the protein level but also at the mRNA level. This also highlights a potentially important role for gigaxonin in RNA regulation under normal and pathological conditions, including in the context of GAN-associated neurodegeneration.

Our study also revealed GAN-associated mis-localization of TDP-43, a neurodegeneration-associated RBP. Pathogenic deposits of TDP-43 within neuronal cytoplasmic inclusions are associated with a number of neurodegenerative disorders, including frontotemporal lobar disease (FTLD) and ALS (De Boer et al., 2021). Major functions of TDP-43 include the regulation of RNA splicing, transport, and stabilization, as well as the formation of stress granules (SGs). SGs are translationally inactive structures containing mRNA, ribosomal binding proteins, and ribosomal components (Wolozin and Ivanov, 2019). In chronic disorders producing persistent stress, SGs may accumulate over time and mature into stable complexes within pathologic tissue — a feature that makes SGs particularly relevant to neurodegenerative diseases. Transcriptomic profiling has shown that *KLHL16/GAN* mRNA is enriched in SGs (Khong et al., 2017), and computational analysis of human *KLHL16* mRNA reveals that it has an exceptionally long 3′ UTR containing numerous RBP binding motifs. The prior observation that ∼40% of *KLHL16/GAN* mRNA is localized to SGs (Khong et al., 2017) is consistent with the generally low level of gigaxonin protein expression and the observation that most SG-enriched mRNAs tend to be long, especially in their 3′ UTR (Khong et al., 2017). By linking GAN to TDP-43 and potentially to SG formation, our findings shed new light on the longstanding morphologic observations of distinctive ‘dense granular bodies’ embedded within NF accumulations in neurons of GAN patients (Asbury et al., 1972). The nature of these dense granular bodies, although consistently cited in ultrastructural evaluations of human GAN, has remained under-investigated and unknown. While the focus in the field has been on gigaxonin as a protein (an E3 ubiquitin ligase adapter), here we shift the focus to other functions of gigaxonin, and specifically to its interaction with RBPs important for neuronal homeostasis.

In conclusion, our data provide the first evidence of RNA/RBP dysfunction in the pathogenesis of GAN-related neurodegeneration. Mechanistic interrogation of altered cell structure and underlying proteostasis dysfunction will be critical for uncovering the pathogenesis of GAN, as well as identifying therapeutic targets for GAN and potentially a wide spectrum of neurodegenerative disorders linked to aberrant accumulations of NFs in neurons (Didonna and Opal, 2019).

## Materials and Methods

### Antibodies

The following primary antibodies and concentrations were utilized: mouse anti-tGFP (Origene, OT12H8, WB: 1:1000, IF: 1:200), rabbit anti-Gigaxonin (Novus, NBP1-49924, WB: 1:500), mouse anti-Gigaxonin (Santa Cruz Biotechnology, F3, WB: 1:200), mouse anti-Actin (Santa Cruz, SPM161, WB: 1:1000), rabbit mouse anti-Pan-actin (Cell Signaling Technology, WB: 1:2000), rabbit anti-TDP-43 (Proteintech, WB: 1:2500, IF: 1:250-500), rabbit anti-PCBP2 (Proteintech, WB: 1:1000), mouse anti-NEFH (Fisher Scientific, RMdO-20, IF: 1:250), chicken anti-phospho-neurofilament-H (Thermo Fisher Scientific, IF: 1:500), and mouse anti-βIII-tubulin (Cell Signaling Technology, TU-20, IF: 1:500). The following secondary antibodies and concentrations were utilized: IRDye 800CW goat anti-rabbit IgG (LI-COR, WB: 1:5000), IRDye 680RD donkey anti-mouse IgG (LI-COR, WB 1:5000), and Alexa 488, 568, 594, and 647-conjugated goat anti-mouse, anti-rabbit, and anti-chicken antibodies (Invitrogen, IF: 1:500).

### Transformation and preparation of pCMV6-AC-GFP vector and wild-type gigaxonin

Transformation for the pCMV6-AC-GFP vector (Origene) and GFP-tagged wild-type gigaxonin construct (pCMV6-AC-GFP vector; Origene) was conducted using XL1-blue supercompetent cells from Aligent Technologies, which were utilized in accordance with product instructions. The cell suspension from the transformation was spread onto an ampicillin agar plate, then incubated overnight at 37°C. Colonies were individually picked and inoculated overnight at 37°C shaking at 225 RPM in LB media (Fisher Scientifiic) with ampicillin (100µg/µL). Plasmid DNA was extracted using the PureLink™ Quick Plasmid Miniprep kit (Invitrogen) and submitted for Sanger sequencing to check for off-target changes in the vector, coding region of gigaxonin, or GFP-tag. After confirming the sequences, a glycerol stock from each previously-inoculated cell suspension was thawed and added 250mL LB media with ampicillin (100µg/µL), then incubated overnight at 37°C shaking at 225 RPM. Plasmid DNA was extracted using a QIAfilter Plasmid Midi Kit (Qiagen) to generate 100ng/µL and 500ng/µL transfection stocks.

### Cell culture and transfection of gigaxonin mutants

HEK293 cells were thawed in MEM (Gibco) with 10% FBS (GenClone, Lot: P121400) and 1% penicillin/streptomycin (pen/strep; Gibco). The medium was changed every 2-3 days and cells were passaged at <95% confluency with 0.05% Trypsin-EDTA (Gibco).

HEK293 cells were plated on 10cm plates at about 40-60% confluency in MEM +10% FBS media (no antibiotics). The following day, the cells were transfected with the pCMV6-AC-GFP vector and gigaxonin wild-type constructs plus Lipofectamine 3000 and p3000 reagents that were utilized in accordance with product instructions (Thermo Fisher Scientific). A Lipofectamine/p3000 only condition was utilized as an untransfected control. Media was changed to MEM +10% FBS + pen/strep media about 5-7 hours after transfection, and cells were harvested for biochemical analysis (see below for details on preparation of protein lysates).

### iPSC generation and differentiation to motor neurons

#### Reprogramming

The protocols for reprogramming GAN patient-derived fibroblasts to induced pluripotent stem cells (iPSCs) and for correcting GAN-iPSCs via CRISPR/Cas9 to generate the isogenic control cell line were described previously (Battaglia et al., 2023).

#### Maintenance

iPSCs were cultured on Matrigel (Corning) with StemFlex Medium (Thermo Fisher Scientific). Media was changed every day and cells were passaged at least once per week with 0.5 mM EDTA dissociation solution.

#### Differentiation

iPSCs were dissociated into single cell suspension with Accutase (Thermo Fisher Scientific) and counted to yield 3×10^5^ cells per well cultured in suspension on an orbital shaker at 37°C with StemScale PSC medium (Thermo Fisher Scientific) supplemented with 10 µM Y27632 (PeproTech). To differentiate the cells to neural progenitor cells (NPCs), the cells were dissociated with Accutase after 48 hours and cultured in PSC neural induction medium (ThermoFisher) supplemented with 10 µM Y27632, 10 µM SB431542 (PeproTech) and 50 nM LDN-193189 (Sigma) to generate neurospheres. At day 7, the cells were cultured with 100 µM Retinoic Acid (Sigma) and 500 nM SAG (Millipore) with daily medium changes and dissociated at 1:3 ratio twice a week until day 12. From day 12 to day 20, the cells were cultured in Neuronal Expansion Medium (1:1 Advanced DMEM/F12 (Thermo Fisher Scientific) and Neural Induction Supplement (Thermo Fisher Scientific) plus 100 µM Retinoic Acid and 500nM SAG). For further differentiation into mature neurons, NPCs were plated on PDL/Laminin-coated plates or coverslips in maturation medium 1 (MM1) for 7 days (1:1 Advanced DMEM/F12 and Neurobasal (Thermo Fisher Scientific); 1X GlutaMAX (Thermo Fisher Scientific), 100mM 2-mercaptoethanol (Thermo Fisher Scientific), 1x B27 supplement (Thermo Fisher Scientific), 0.5x N2 supplement (Thermo Fisher Scientific), 1x NEAA (Thermo Fisher Scientific) and 2.5 g/ml insulin (Sigma-Aldrich)) supplemented with 10 ng/ml BDNF (Thermo Fisher Scientific), 10 ng/ml GDNF (Thermo Fisher Scientific), 10 µM Forskolin (Stem Cell Technologies), 100 µM IBMX (Tocris), and 10 µM DAPT (R&D). Cells were cultured at least two weeks to achieve maturation.

#### Differentiation on microfluidic chips

Using the differentiation protocol described above, iPSC-derived motor neurons were cultured on microfluidic chips with distinct soma and axonal compartments (Xona Microfluidics®; XC900). Each chip was coated with poly-L-Ornithine for 16 hours, followed by laminin for 2-3 hours, followed by washes and media conditioning according to the manufacturer’s protocol. NPCs were dissociated to small clusters and ∼120,000 cells were seeded onto each chip, followed by media changes every two days. The cells were cultured for 30 days to achieve maturation in the microfluidic chips.

### Preparation of protein lysates and immunoblotting

*GFP-tagged gigaxonin transfection lysates:* For biochemical analysis, transfection lysates were prepared by lysing cells from each condition in 1mL cold TritonX-100 buffer (1% TritonX-100, 0.5 M EDTA, PBS, ddH2O) with protease/phosphatase inhibitors (Halt™ Protease & Phosphatase Inhibitor Cocktail, EDTA-free, 100X; Thermo Fisher Scientific). The lysates were spun down in a tabletop centrifuge at 15,000 RPM for 10 minutes at 4°C to separate the TritonX-100 detergent soluble and insoluble fractions; the supernatant for each sample was transferred to a new tube (detergent soluble fraction). Total protein concentration for the detergent soluble fractions were measured with Pierce™ BCA Protein Assay Kit (Thermo Fisher Scientific) and Biotek Synergy HT plate reader according to manufacturer’s instructions; total protein concentrations were calculated from the recorded absorbance values. The concentrations were used to calculate 1µg/µL stocks for immunoblotting and immunoprecipitation (see below for more details on immunoprecipitation); the samples were diluted with TritonX-100 buffer (+protease/phosphatase inhibitors). For immunoblotting, hot 2x SDS sample buffer (Thermo Fisher Scientific) reduced with 5% 2-mercaptoethanol (Sigma) was added 1:1 to 1µg/µL samples for each condition, then samples were heated at 95°C for 5 minutes.

#### GAN patient iPSC-NPC and iPSC-MN lysates

Following differentiation and maturation, iPSC-neural progenitor cells (NPCs) and iPSC-motor neurons (3x clones G332R point mutant line, 3x clones isogenic control line) were washed with HBSS, collected and pelleted, and flash frozen as cell pellets. The cell pellets were thawed on ice, resuspended in with TritonX-100 buffer (1% Triton X-100, 0.5 M EDTA, PBS, ddH2O) with protease/phosphatase inhibitors (cOmplete™ Protease Inhibitor Cocktail (Roche), PhosSTOP phosphatase inhibitor cocktail (Roche)) to lyse the cells. Total cell lysates were prepared by transferring a fraction of the lysate to a new tube, adding an equal volume of hot sample buffer reduced with 5% 2-mercaptoethanol (Sigma), and heating at 95°C for 5 minutes.

#### Immunoblotting

Protein lysates were loaded with equal volumes and separated on 10% or 4-20% Novex™ WedgeWell™ Tris-Glycine gels (Thermo Fisher Scientific) for 35 minutes at 225V or 1 hour 20-40 minutes at 120V, then transferred for 2 hours at 80V at 4°C or overnight at 40V at 4°C onto nitrocellulose membranes. Gels were stained with Coomassie and destained following each transfer to verify normalization. The membranes were blocked in 5% non-fat milk (NFM) dissolved into 0.1% tween 20/PBS (PBST) at room temperature for 30-60 minutes. The membranes were incubated in primary antibodies diluted in 5% NFM/PBST for 1 hour at room temperature or overnight at 4°C (see concentrations above), then washed 3x with PBST for 5 minutes each. The membranes were incubated with secondary antibodies diluted in 5% NFM/PBST at room temperature for 1 hour (see concentrations above), washed 3x with PBST and 1x with PBS for 5 minutes each, and then scanned with a LI-COR Odyssey CLx machine. Image Studio version 5.2 (LI-COR) was used to perform densitometry on immunoblots, and Adobe Photoshop was used for densitometry on gels stained with Coomassie. Pan-actin bands on membranes and total protein concentration on gels were quantified and utilized to normalize loading volumes for the samples as needed before blotting for proteins of interest.

### Immunoprecipitation with anti-GFP magnetic beads

ChromoTek TurboGFP-Trap Magnetic Agarose beads (Proteintech) were utilized for immunoprecipitation (IP) of GFP-tagged vector and gigaxonin wild-type constructs in accordance with product instructions. In brief, the beads were equilibrated with 3x washes with TritonX-100 buffer (1% Triton X-100, 0.5 M EDTA, PBS, ddH2O) with protease/phosphatase inhibitors (Halt™ Protease & Phosphatase Inhibitor Cocktail, EDTA-free, 100X; Thermo Fisher Scientific); washes were manually removed when beads were separated using a magnetic rack. Following equilibration, detergent soluble 1µg/µL stocks were loaded onto magnetic beads and incubated overnight at 4°C while rotating. Following incubation, the captured protein was eluted in 50mM Ammonium Bicarbonate (pH 7.8) for mass spectrometry-based proteomics analysis and immunoblotting.

### Immunofluorescence staining, imaging, and analysis

GAN patient iPSC-NPCs and iPSC-MNs were briefly rinsed 1-2x with PBS, then fixed in 4% paraformaldehyde in PBS (Thermo Fisher Scientific) for 15-30 minutes at room temperature and followed by 3x 5-minute PBS washes. The cells were then permeabilized for 5 minutes with 0.1% Triton X-100 in PBS, washed 3x with for 5 minutes each, and incubated with blocking buffer for 1 hour at room temperature. After blocking, the cells were incubated overnight with primary antibodies at 4°C (see concentrations above), followed by 3x 5 min PBS washes, then incubated with Alexa Fluor-conjugated secondary antibodies (see concentrations above) at room temperature for 1 hour and washed 3x with PBS for 5 minutes each. Both primary and secondary antibodies were diluted in blocking buffer. Finally, cells were incubated with DAPI (Invitrogen), washed 2-3x with PBS for 5 minutes each, and mounted in Fluoromount-G (SouthernBiotech) overnight. iPSC-motor neurons cultured in the microfluidic chips were stained using the same procedure as above. SlowFade® anti-fade reagent (Thermo Fisher Scientific) was added to microfluidic chambers to minimize signal quenching. Images were acquired on a Zeiss 880 confocal laser scanning microscope using 40x or 63x oil immersion objectives, or acquired on Olympus FluoView FV1000 confocal microscope using a 30x silicon immersion objective.

### Electron microscopy

GAN patient iPSCs were differentiated to motor neurons as described above, fixed in 2.5% glutaraldehyde in 0.1M sodium cacodylate buffer (pH 7.4) for one hour at room temperature, and stored at 4°C. The cells were washed 3 times in 0.1M sodium cacodylate buffer followed by post-fixation in 1% buffered osmium tetroxide for 1 hour. After three washes in deionized water, the cells were dehydrated in ethanol, infiltrated and embedded in situ in PolyBed 812 epoxy resin (Polysciences, Inc, Warrington, PA). The cell monolayer was sectioned *en face* to the substrate with a diamond knife and Leica UCT Ultramicrotome (Leica Microsystems, Inc, Buffalo Grove, IL). Ultrathin sections (70 nm) were mounted on 200 mesh copper grids and stained with 4% uranyl acetate and lead citrate. The sections were observed and digital images were taken using a JEOL JEM-1230 transmission electron microscope operating at 80kV (JEOL USA, Inc, Peabody, MA) equipped with a Gatan Orius SC1000 CCD Digital Camera (Gatan, Inc, Pleasanton, CA).

### RNAscope in situ hybridization

The RNAscope® Multiplex Fluorescent Assay protocol from Advanced Cell Diagnostics (ACDBio; 323100) was utilized to probe for *KLHL16* mRNA expression as described previously (Battaglia et al., 2023).

### Mass spectrometry-based proteomics analysis

#### GFP-tagged gigaxonin immunoprecipitation

The immunoprecipitation samples were prepared as described above and stored in 50mM Ammonium Bicarbonate (pH 7.8), then submitted for mass spectrometry-based proteomic analysis. In brief, on-bead tryptic digestion was performed, followed by peptide extraction and C18 desalting cleaning. Each sample was analyzed by LC-MS/MS using the Thermo Easy nLC 1200-QExactive HF. The data were processed using MaxQuant (v1.6.15.0) – the data were searched against the Uniprot Human database (∼20,000 sequences), a common contaminants database (∼250 sequences), and gigaxonin sequences. The MaxQuant results were filtered and analyzed in Perseus, and reverse hits and proteins with 1 peptide (single hits) were removed. Perseus was used for imputation, log2 fold change and p-value calculations.

#### GAN neural progenitor cells (NPCs)

Following differentiation and maturation, iPSC-neural progenitor cells (NPCs; 3x clones G332R point mutant line, 3x clones isogenic control line) were washed with HBSS, collected and pelleted, and flash frozen as cell pellets. Total cell lysates were prepared as described above and normalized volumes for each lysate were loaded into NuPAGE™ 4 to 12% gels (Thermo Fisher Scientific). The gels were run at 120V until the samples entered the gels and were stacked about 1cm. The gels were removed and washed 5x for 5 minutes with molecular grade water to remove excess sample buffer. Gels were incubated with GelCode Blue Stain Reagent for 1 hour at room temperature on a shaker and periodically monitored for protein band development. The gels were destained by incubating with molecular grade water at room temperature for 1-2 hours on a shaker and changing water multiple times throughout this time period. The band for each sample was excised from the gel and stored in molecular grade water, then the bands were submitted for mass spectrometry-based proteomic analysis. In brief, in-gel trypsin digestion was performed, followed by peptide extraction and desalting using Pierce peptide desalting spin columns. The peptides were quantified using Pierce BCA fluorometric peptide quantitation assay. After normalizing the sample concentrations, the samples were analyzed by LC-MS/MS using a Thermo Ultimate 3000-Exploris480. MaxQuant was used to identify and quantify proteins; the data were searched against a Uniprot Human database (∼20,000 proteins) and appended with a common contaminants database. The data were imported into Argonaut for normalization, filtering, imputation, and visualization.

### Data Analysis and Statistics

The proteomics data in Figures 1B, 1E, 1F, and 2D were analyzed using DAVID (Sherman et al., 2022) bioinformatics resource. Bar graphs and heat maps were generated with the GraphPad Prism software and statistical analysis was performed via unpaired t-test or one-way ANOVA, as denoted in the figure legends. Volcano plot of the mass spec results (Fig. 1D) was generated using the Volcanoser web app(Goedhart and Luijsterburg, 2020).

## Summary of Supplemental Material

- **Supplemental Table 1:** List of all soluble gigaxonin interactors
- **Supplemental Table 2:** List of gigaxonin RBP interactors
- **Supplemental Table 3:** List of differentially expressed proteins in GAN vs Control NPCs.

## Acknowledgments

The authors thank Dr. Laura Herring for assistance with mass spectrometry analysis, Dr. Adriana Beltran for assistance with GAN iPSC derivation and differentiation, and Drs. Anne Taylor and Tharkika Nagendran for assistance with the use of Xona microfluidic chips. This study was funded by Hannah’s Hope Fund and NIH grants R21NS121578 (NTS, DA), GM122741 and AHA grant 24PRE1193707 (training awards to C.P.), F99AG068523 (RAB), R01NS127204 (PO), and R01NS082351 (PO).

## Notes

### Competing Interest Statement

The authors have declared no competing interest.

